# DOT1L inhibition reveals a distinct class of enhancers dependent on H3K79 methylation

**DOI:** 10.1101/383489

**Authors:** Laura Godfrey, Nicholas T. Crump, Ross Thorne, I-Jun Lau, Emmanouela Repapi, Dimitra Dimou, Jelena M. Telenius, A. Marieke Oudelaar, Damien J. Downes, Paresh Vyas, Jim R. Hughes, Thomas A. Milne

## Abstract

Enhancer elements are a key regulatory feature of many important genes. Several general features including the presence of specific histone modifications are used to identify and subcategorize enhancers. Here we identify a distinct subset of enhancers in leukemia cells that are functionally dependent upon H3K79me3. Using the DOT1L inhibitor, EPZ-5676, we show that loss of H3K79me3 at these H3K79me3 enhancer elements (KEEs) leads to reduced chromatin accessibility, histone acetylation and transcription factor binding. We then use Capture-C, a high-resolution chromosome conformation capture technique, to show that H3K79me3 is required for KEE interactions with the promoter as well as transcription of the associated genes. Together these data implicate H3K79me3 in having a functional role at a subset of active enhancers where it helps maintain histone acetylation and chromatin accessibility, potentially by promoting phase-separated condensates.

## Introduction

Enhancers are key regulatory elements that contribute to gene expression. They function in part by acting as docking sites for transcription factors, potentially resulting in the accumulation of protein complexes that create phase separated condensates generating distinct regulatory domains (Cho et al., 2018; Sabari et al., 2018). In general, active enhancers display characteristics such as an open chromatin conformation and post translational histone modifications such as H3 lysine 4 monomethylation (H3K4me1) and H3 lysine 27 acetylation (H3K27ac) (Creyghton et al., 2010; Heintzman and Ren, 2009; Heintzman et al., 2007; Rada-Iglesias et al., 2011). These features can be used to help identify potentially functional enhancers genome wide in different cell types. Attempts to use other unique features to subclassify enhancers into different functional groups has resulted in the identification of super-enhancers (Hnisz et al., 2013; Loven et al., 2013; Whyte et al., 2013). It has also been recognized that H3K79me3 is an important marker of some developmentally active enhancers (Bonn et al., 2012; Gilan et al., 2016; Markenscoff-Papadimitriou et al., 2014), but the functional significance of H3K79me3 at enhancers has not been established.

Disruptor of telomeric silencing 1-like (DOT1L) is the only known methyltransferase for histone H3 lysine 79 di or trimethylation (H3K79me2/3) (Feng et al., 2002). H3K79me2/3 is mainly found within the body of active genes and is canonically associated with transcription elongation (Biswas et al., 2011; Feng et al., 2002; Mohan et al., 2010; Mueller et al., 2007; Steger et al., 2008). The precise role of H3K79me2/3 in promoting transcription elongation is unknown, but one model suggests that H3K79me2/3 functions in part by inhibiting histone deacetylase activity and by preventing the formation of H3 lysine 9 trimethyl (H3K9me3) repressive domains (Chen et al., 2015). Some additional support for this model comes from recent evidence that KDM2B may act as a histone demethylase for H3K79me2/3 leading to transcriptional silencing by recruitment of SIRT1 and gain of H3K9me3 (Kang et al., 2018). H3K79me2/3 has also been shown to be important in human disease. In particular, H3K79me2/3 is an important driver of leukemogenesis, mainly in a rare subset of leukemias caused by rearrangements of the *Mixed lineage leukemia* gene (MLL-r) (Bernt et al., 2011; Deshpande et al., 2013; Krivtsov et al., 2008; Milne et al., 2005; Mueller et al., 2007; Okada et al., 2005). The most common MLL rearrangements are chromosome translocations that fuse MLL in-frame with over 120 other genes creating novel fusion proteins, although MLL-AF4 is the most common MLL fusion produced (Meyer et al., 2013). MLL-AF4 is a major cause of incurable acute lymphoblastic leukemia (ALL) in infants and children (Andersson et al., 2012; Meyer et al., 2013; Milne, 2017), and increased H3K79me2/3 correlates with increased transcription of MLL-AF4 bound genes (Benito et al., 2015; Bernt et al., 2011; Godfrey et al., 2017; Kerry et al., 2017; Lin et al., 2016; Wilkinson et al., 2013), likely due to aberrant recruitment of DOT1L, (Biswas et al., 2011; Gilan et al., 2016; Milne et al., 2005; Yokoyama et al., 2010). The highly specific DOT1L inhibitor, EPZ-5676 (pinometostat), has moderate clinical activity in patients (Stein et al., 2018) and has been extensively characterised in numerous studies as an excellent tool to study the molecular function of DOT1L (Chen et al., 2015; Daigle et al., 2013).

In this study, we perform a genome wide analysis in MLL-AF4 human leukemia cells (SEM) and discover a set of enhancers that are marked with H3K79me3 which we term H3<u>K</u>79me3 <u>E</u>nhancer <u>E</u>lements (KEEs). We go on to demonstrate that H3K79me3 plays a functional role at these enhancers with loss of H3K79me3 leading to a specific reduction of H3K27ac, chromatin accessibility and transcription factor binding at KEEs. Strikingly, we observe that this perturbation in enhancer function leads to a disruption in chromatin interactions between KEEs and the promoter of the regulated gene. This is coupled with a downregulation of transcription. Together, these results define a distinct class of enhancers which are dependent on H3K79me3 to maintain an active, open chromatin configuration and sustain interactions with gene promoters.

## Results

### H3K79me3 marks a new class of enhancers in leukemia

Whilst H3K79me3 is canonically associated with active gene bodies due to its association with transcription elongation (Mueller et al., 2007; Steger et al., 2008), the role of H3K79me3 in enhancer function has only been partially explored (Bonn et al., 2012; Gilan et al., 2016; Markenscoff-Papadimitriou et al., 2014). To identify H3K79me3-marked enhancers genome wide in leukemia cells, we used the automated chromatin-state discovery package ChromHMM (Ernst and Kellis, 2012, 2017) and classified enhancers based on the presence of H3K27ac and H3K4me1 (Figure 1A). We identified 16877 putative enhancers, of which 3621 (21%) were directly marked with H3K79me3 (Figure 1B), indicating that this modification may play a role at a significant subset of enhancers. Notably, the vast majority of these enhancers (3502) were intragenic (Figure 1B, S1A). This fits with past analyses that suggest that H3K79me3 is enriched in the body of genes (Bernt et al., 2011; Kerry et al., 2017; Steger et al., 2008) compared to intergenic regions, but also highlights a class of enhancers which are marked with H3K79me3. Enhancer lengths were comparable, with median lengths of 1 kb and 0.8 kb for H3K79me3-marked and unmarked enhancers, respectively (Figure S1E). For convenience, we refer to H3<u>K</u>79me3 marked <u>e</u>nhancer <u>e</u>lements as “KEEs” and non-H3K79me marked enhancer elements as “non-KEEs”.

**Figure 1.**
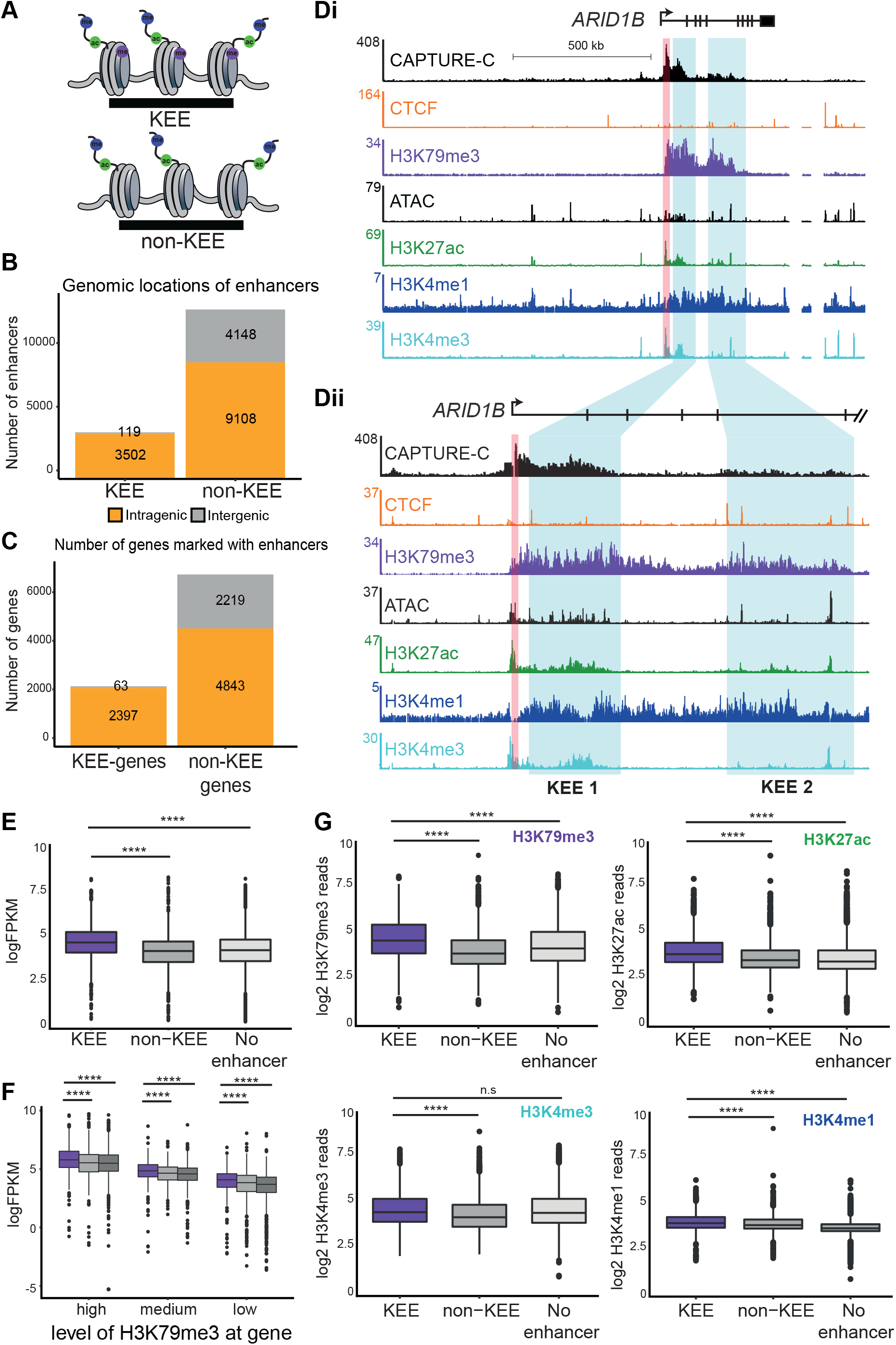
H3K79me3 marks a new class of enhancers in leukemia (**A**) Schematic diagram demonstrating how KEEs were categorized (**B**) Genomic location (intragenic = orange and intergenic = grey) of KEEs and non-KEEs. Enhancers identified using ChromHMM based upon H3K27ac, H3K4me1 and H3K79me3 ChIP seq peaks in SEM cells. (**C**) Number of genes associated with a KEE or non-KEE (intragenic = orange, intergenic = grey). Enhancers paired with the nearest gene (**D**) Capture-C, ChIP seq and ATAC seq at *ARID1B* performed in SEM cells. Blue bars indicate KEE clusters 1 and 2 (**E**) Average expression of genes (logFPKM) associated with a KEE (purple), non-KEE (grey) and no enhancer (light grey) (*** =p-value <0.0001 using a Mann-Whitney *U* test). (**F**) Average expression (logFPKM) of genes associated with a KEE (purple), non-KEE (grey) and no enhancer (light grey) categorised into high, medium, low levels of H3K79me found within the gene body. (**** =p-value <0.0001 using a Mann Whitney U test). (**G**) Level of H3K79me3 (purple), H3K27ac (green), H3K4me1 (teal) and H3K4me1 (blue) at genes associated with KEEs (purple), non-KEEs (grey) and no enhancer (light grey), normalised for gene length. (**** = p-value <0.0001, n.s. = non-significant, Mann-Whitney *U* test). See also Figure S1.

To understand the functional significance of KEEs, we first wanted to determine how many genes were potentially regulated by a KEE. We used proximity to assign KEEs and non-KEEs to the nearest gene. We found that although most genes were associated exclusively with non-KEEs (non-KEE gene), 2460 genes were associated with at least one KEE (KEE-genes) (Figure 1C), suggesting KEEs may directly regulate the expression of a subset of genes. To validate this observation, we performed high resolution chromosome conformation capture at a subset of key target genes (See Table S2) using next generation Capture-C (Davies et al., 2016), which is capable of producing high resolution interaction profiles of loci of interest (see also Figure 6). As an example, the *ARID1B* gene contains two intragenic KEE clusters, marked with H3K79me3 as well as H3K4me1 and H3K27ac and open chromatin as demonstrated by ATAC seq (Figure 1 Di and Dii, shaded regions). Strikingly, Capture-C from the viewpoint of the *ARID1B* promoter revealed interactions with both KEE clusters, suggesting they might directly regulate *ARID1B* expression (Figure 1 Di and Dii). Conversely, the SPI1 promoter interacts with both KEEs and non-KEEs (Figure S1B). However, Capture-C shows that the major region of interaction is with an upstream non-KEE cluster (shaded) marked by multiple peaks of H3K4me1, H3K27ac and ATAC-seq, which is centered around a previously identified SPI1 upstream response element (URE) (Huang et al., 2008), suggesting that this is the major regulatory element.

We next investigated the effect of KEEs on gene transcription by nascent RNA-seq. KEE genes have significantly higher levels of expression compared to non-KEE genes and those with no assigned enhancer (Figure 1E, Table S3-S4). Since genes without H3K79me3 generally have lower expression (Bernt et al., 2011; Kerry et al., 2017; Steger et al., 2008), and many non-KEE and non-enhancer assigned genes lack H3K79me3 in the gene body (Figure S1C-D), we excluded non-H3K79me3 genes from our analysis to avoid this bias. Interestingly, even with non-H3K79me3 marked genes removed, KEE genes have higher levels of expression compared to non-KEE genes and those with no assigned enhancer (Figure 1E). Notably, KEE-genes have higher expression levels even when comparing genes with similar levels of H3K79me3 in the gene body, suggesting that the enhancers may be increasing transcription rates (Figure 1F). Consistent with this observation, although there were no differences in H3K4me3 levels, we found higher levels of active histone modifications (H3K27ac and H3K4me1) at KEE genes compared to non-KEE genes (Figure 1G).

Previous work to identify novel classes of enhancers established the concept of superenhancers, marked by MED13, BRD4, H3K27ac and H3K4me1 (Hnisz et al., 2013; Loven et al., 2013; Whyte et al., 2013). It was recently shown that H3K79me3 may have a role at a subset of super-enhancers (Gilan et al., 2016), so we wanted to determine whether KEEs were distinct from super-enhancers. We identified super-enhancers in MLL-AF4 cells as in the original study and found that most KEEs are not super-enhancers (Figure S1H), and there is no specific enrichment for KEEs among super-enhancers. This suggests that KEEs, in MLL-AF4 cells, are distinct from super-enhancers.

We next wanted to determine if KEEs are a distinct category of enhancer and whether they are associated with any documented regulatory features in leukemia cells. Recent work from our lab has shown that a subset of key MLL-AF4 gene targets associated with a poor prognosis are distinguished by high levels of gene expression as well as broad binding of MLL-AF4 across the gene and high levels of H3K79me3. These genes were termed MLL-AF4 spreading targets (Kerry et al., 2017). We therefore asked whether the presence of H3K79me3 at KEEs is a consequence of MLL-AF4 spreading into intragenic regions. Interestingly, whilst MLL-AF4 spreading targets are enriched for KEEs compared to nonspreading targets (Figure S1F-G), suggesting there may be a link between these two features, the vast majority of KEE genes are not spreading targets. However, a role for MLL-AF4 in promoting KEEs at a subset of genes could be a key contributor to leukemia progression. Taken together, these data suggest that KEEs are a distinct category of enhancers which are not limited to super-enhancers or MLL-AF4 spreading targets.

### Loss of H3K79me at enhancers leads to a reduction in transcription at KEE-genes

Since KEEs are associated with highly expressed genes, we wanted to better understand the function of H3K79me3 at KEEs in gene regulation. To accomplish this, we used the DOT1L small molecule inhibitor EPZ-5676 (Daigle et al., 2013) to diminish H3K79me3 genome-wide (Figure 2A). Treatment with 2 μM of EPZ-5676 led to a near-complete loss of global H3K79me3 levels as measured by ChIP-rx (Orlando et al., 2014) and western blot (Figure 2B, S2A-2B). We performed nascent RNA seq to identify genes dependent on H3K79me3 for transcription and identified 2601 up- and 2462 downregulated genes (Figure 2C, S2D-F) following DOT1L inhibition (DOT1Li). Downregulated genes were more likely to be marked by H3K79me3 (Figure 2D, S2C-D), consistent with a role in active transcription (Steger et al., 2008). For example, at the KEE gene *ARID1B*, loss of H3K79me3 is coupled with downregulation of transcription (Figure 2E) and RT-PCR validates loss of expression at several other KEE genes (Figure S2E). *BCL2*, which is primarily regulated by a downstream non-KEE (Godfrey et al., 2017) but contains high levels of H3K79me3 in the gene body, also demonstrated a reduction in transcription following DOT1Li (Figure S2F), highlighting the potential for non-enhancer roles in transcription for H3K79me3 at some non-KEE genes. However, we also observed non-KEE genes which were marked with H3K79me3 in the gene body but were not sensitive to DOT1Li, such as *FLT3* (Figure 2D, S2F).

**Figure 2.**
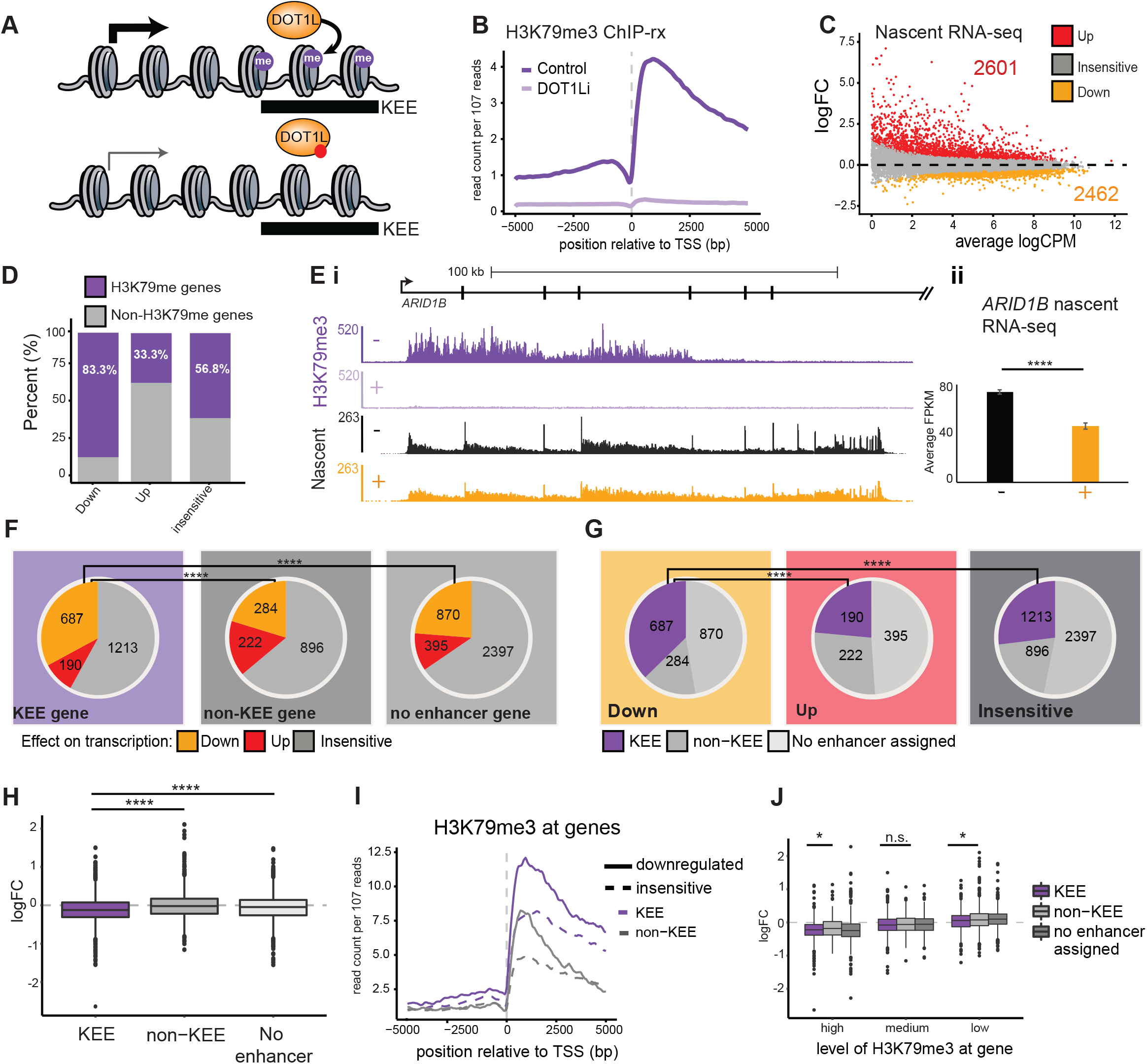
Loss of H3K79me at enhancers leads to a reduction in transcription at KEE-genes. (**A**) Schematic showing reduction of gene expression following loss of H3K79me at KEE genes. The red dot is meant to indicate the presence of the inhibitor EPZ-5676 (**B**) Metagene plot of H3K79me3 ChIP-rx signal following DOT1Li (EPZ-5676) treatment (+, lilac) compared to control (-, DMSO) treatment (purple) in SEM cells (**C**) MA plot of nascent RNA sequencing data showing differential gene expression in SEM cells treated with DOT1Li compared to control. Differential expression = FDR <0.05 (**D**) Proportion of differentially expressed genes and insensitive genes which are directly marked with H3K79me3 (**E**) (**i**) H3K79me3 ChIP-rx tracks showing control (-, purple) and DOT1Li (+, lilac) and nascent RNA sequencing tracks showing control (-, black) and DOT1Li (+, orange) at *ARID1B*. (**** = FDR <0.0001) (ii) Bar chart displaying average FPKM at ARID1B in control (-, black) and DOT1Li (+, orange) conditions. P-value (FDR) <0.0001 (**F**) Proportion of KEE-genes, non-KEE genes and no enhancer genes which are upregulated (red), downregulated (orange) or insensitive (grey) following DOT1Li (****=p-value <0.0001, Fishers exact test). (**G**) Proportion of upregulated, downregulated or insensitive genes which are associated with a KEE (purple), a non-KEE grey) or no enhancer (light grey) (****=p-value <0.0001, Fishers exact test). (**H**) Average logFC of KEE-genes (purple), non-KEE genes (grey) and no enhancer genes (light grey) (**I**) Level of H3K79me3 found at downregulated and insensitive genes which are associated with a KEE or non-KEE. (**** = p-value <0.0001, n.s. = non-significant, Mann-Whitney *U* test) (**J**) Average logFC of KEE-genes (purple), non-KEE genes (grey) and no enhancer genes (light grey) when categorised into high, medium or low levels of H3K79me3 found in the gene body, normalised for gene length. (* = p-value <0.05, n.s. = non-significant, Mann-Whitney *U* test) See also Figure S2.

KEE genes were significantly more sensitive to DOT1Li, with 33% (687) downregulated, compared to 20% (284) of non-KEE genes and 23% (870) of no enhancer assigned genes (Figure 2F). Furthermore, 37% of downregulated genes were KEE genes and demonstrated an increased sensitivity to DOT1Li (Figure 2G-H). A meta-analysis across different gene target sets indicates that H3K79me3 levels are highest at KEE genes compared to non-KEEs genes and genes with no enhancer (Figure 2I). To better understand whether high levels of H3K79me3 are predictive of sensitivity to DOT1Li, we grouped KEE genes, non-KEE genes and genes with no enhancer by high, medium and low levels of H3K79me3 (Figure 2J). We observed that high levels of H3K79me3 in general lead to an increase in differential expression, however, within groups, KEE genes were generally more sensitive to loss of H3K79me3 than non-KEE genes (Figure 2J). Taken together, these results show that a subset of genes associated with a KEE display reduced transcription following DOT1Li, implicating H3K79me3 in enhancer function.

### Loss of H3K79me leads to a reduction in chromatin accessibility at KEEs

Our results so far indicate that H3K79me3 has a specific role in transcription of a subset of KEE genes (Figure 2A). This suggests that these genes are regulated by the KEE and that loss of H3K79me3 directly impacts enhancer function. Active enhancers are associated with regions of accessible chromatin as well as H3K27ac and H3K4me1 (Creyghton et al., 2010; Heintzman and Ren, 2009; Heintzman et al., 2007; Rada-Iglesias et al., 2011). If H3K79me3 regulates the activity of KEEs, then we would expect DOT1Li to produce a reduction in active enhancer features such as chromatin accessibility (Figure 3A). Changes in accessibility are not only reflective of changes in chromatin structure but can also serve as a convenient proxy for changes in transcription factor binding at both enhancers and promoters (Buenrostro et al., 2013). We used ATAC-seq following DOT1Li to determine whether loss of H3K79me3 leads to a reduction in chromatin accessibility at enhancers. In general, we noticed that decreased ATAC-seq peaks were associated with higher levels of H3K79me3 and downregulation of transcription following DOT1Li (Figure S3A-C, Table S5). Although many ATAC peaks did not change in response to DOT1Li, we observed decreases and increases in chromatin accessibility at both KEE-genes and non-KEE-genes, with enrichment of decreased peaks at KEE-genes (Figure 3B-C, S3D, Table S5).

**Figure 3.**
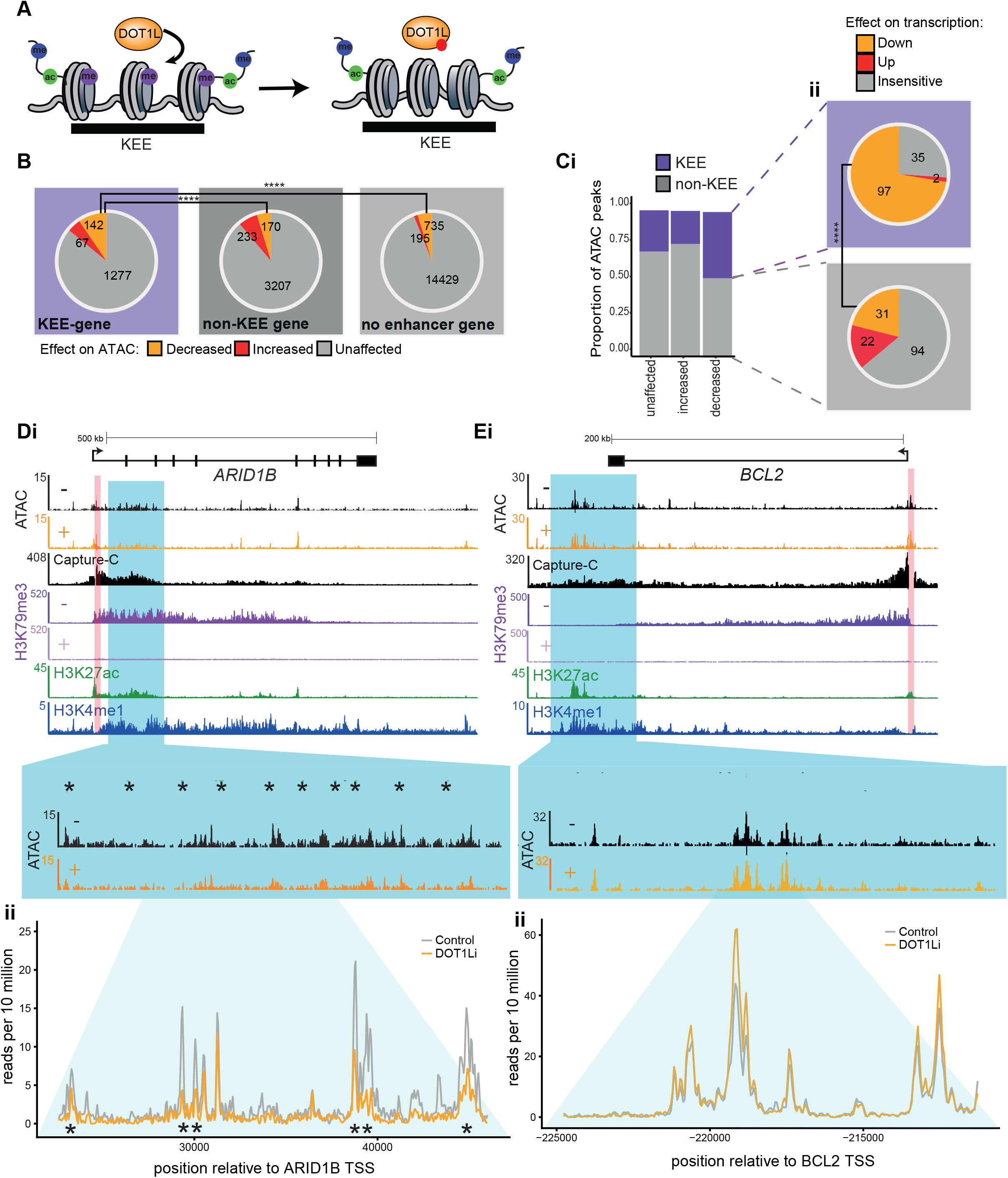
Loss of H3K79me3 leads to reduction in chromatin accessibility at H3K79me3 enhancers (**A**) Schematic demonstrating a loss of chromatin accessibility at KEEs following DOT1Li. (**B**) Proportion of ATAC peaks within KEE genes, non-KEE genes and genes with no enhancer that show increases (red), decreases (orange) or no change (grey) following DOT1Li (<***=p-value <0.0001, Fishers exact test). (**C**)(i) Proportion of increased, decreased or no change in ATAC found with a KEE (purple) or non-KEE (grey) (ii) Proportion of decreased ATAC peaks that are associated with transcriptionally downregulated (orange), upregulated (red) or insensitive (grey) KEE (purple) or non-KEE gene (grey) (****=p-value <0.0001, Fishers exact test). (Di) ATAC seq for *ARID1B* in control (-, black) and DOT1Li (+, orange). Blue bars represent KEE region and zoomed in regions. Stars represent significantly reduced ATAC peaks. (ii) Overlay between ATAC signal at *ARID1B* KEE1 in control (black) and DOT1Li (orange). (Ei) ATAC seq for *BCL2* in control (-, black) and DOT1Li (+, orange) Blue bars represent non-KEE region and zoomed in regions (ii) Overlay between ATAC signal at *BCL2* non-KEE1 in control (black) and DOT1Li (orange). See also Figure S3.

To understand whether these changes in chromatin accessibility were relevant to enhancer function, we compared decreased ATAC peaks at KEEs or non-KEEs to changes in transcription of the associated genes. Strikingly, 73% of decreased ATAC peaks within KEEs correlated with downregulation of transcription of the corresponding genes, in contrast to only 21% of decreased peaks at non-KEEs (Figure 3C). This suggests that reduced chromatin accessibility at KEEs, in response to loss of H3K79me3, disrupts enhancer function and subsequent activation of KEE-genes. This is exemplified by *ARID1B*, which contains two KEE clusters and demonstrates a reduction in chromatin accessibility and transcription (Figure 3D). In contrast, at *BCL2* although loss of H3K79me3 reduces transcription (Figure S2F, S3E) (Benito et al., 2015; Godfrey et al., 2017), there are no chromatin accessibility changes observed in the non-KEE at the 3’ end of the gene (Figure 3E). This suggests that KEEs are dependent upon H3K79me3 for maintenance of open chromatin, a feature which marks active enhancers.

### Loss of H3K79me leads to a disruption of H3K27ac at KEEs

Given that we observed a reduction in chromatin accessibility at KEEs, we hypothesised that H3K27ac and H3K4me1, histone modifications associated with active enhancers, might also decrease upon loss of H3K79me3 (Figure 4A). Strikingly, H3K27ac displayed a clear genome-wide decrease exclusively at KEEs but not at non-KEEs (Figure 4B-C), despite the fact that there were no global changes in H3K27ac (Figure S4A).

**Figure 4.**
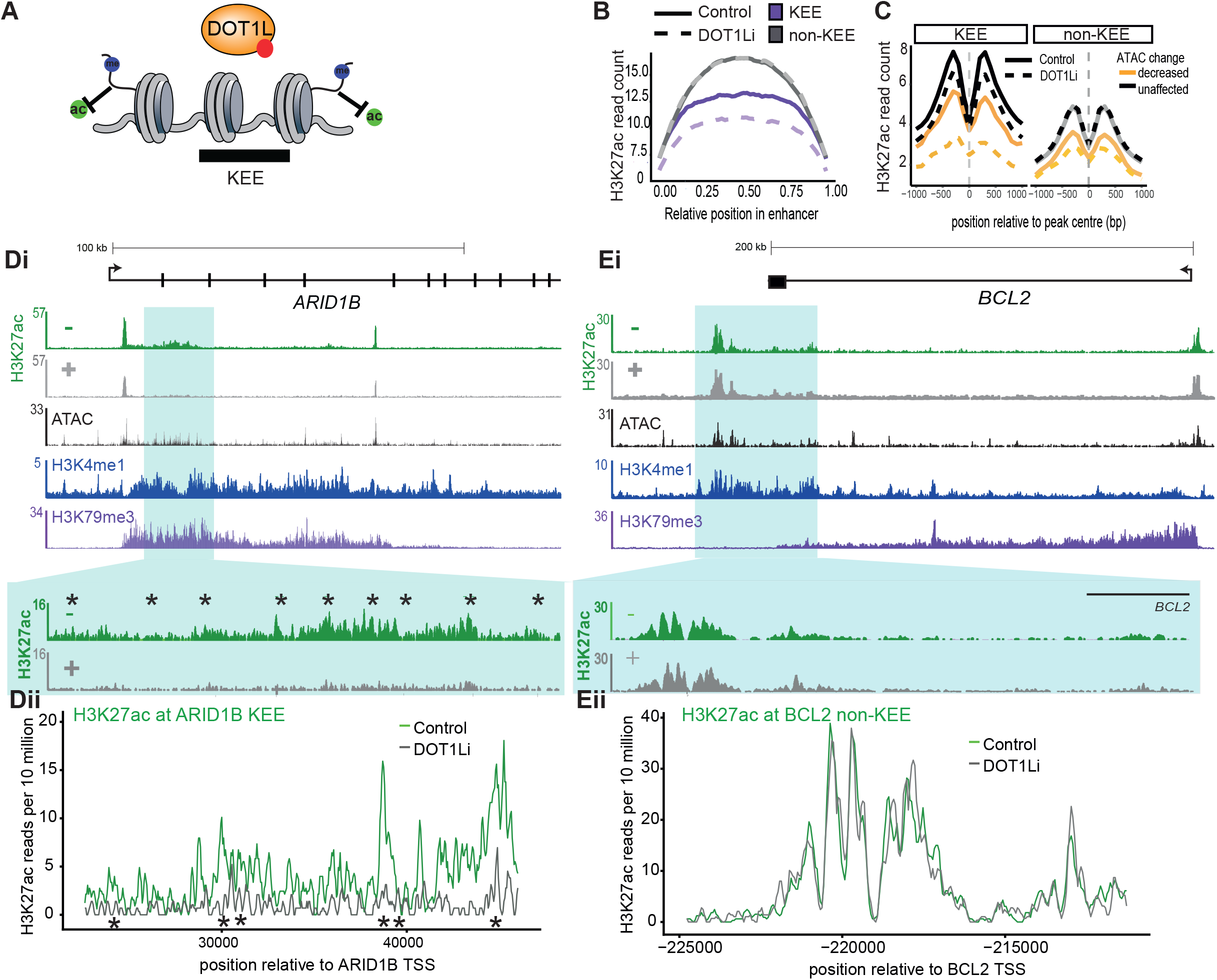
Loss of H3K79me3 and chromatin accessibility perturbs histone acetylation at H3K79me3 enhancers (**A**) Schematic representing a loss of H3K27ac at KEEs following DOT1Li. (**B**) Metaplot of H3K27ac ChIP seq signal centered at KEEs (purple) vs non-KEEs (grey) in control (full line) and DOT1Li (dashed line). (**C**) Metaplot of H3K27ac levels centered at ATAC peaks within KEEs and non-KEEs. Decreased ATAC peaks shown in orange and unaffected in black in control condition (full line) and DOT1Li (dashed line). (Di) H3K27ac ChIP seq at *ARID1B* in control (-, green) and DOT1Li (+, light green). Blue bars highlight KEE regions. Stars represent significant reductions in ATAC signal (ii) Graph depicts overlay between H3K27ac signal at *ARID1B* KEE1 in control (green) and DOT1Li (light green). (Ei) H3K27ac ChIP seq at *BCL2* in control (-, green) and DOT1Li (+, light green). Blue bars highlight KEE regions. Stars represent significant reductions in ATAC signal (ii) Overlay between H3K27ac signal at *BCL2* non-KEE in control (green) and DOT1Li (light green) conditions. See also Figure S4.

These genome wide results can be clearly observed at the *ARID1B* KEEs where loss of H3K79me3 results in a significant loss of H3K27ac and concomitant with chromatin accessibility (Figure 4D). Reductions in H3K9ac were also observed at the *ARID1B* KEEs (Figure S4B), suggesting that there may be a more general loss of histone acetylation marks. This contrasts strongly with the non-KEE of *BCL2* where there is no change in H3K27ac (Figure 4E), despite that fact that *BCL2* is mildly downregulated following DOT1Li (Figure S3E). Interestingly, KEEs that display reduced ATAC and H3K27ac signal have generally lower levels of H3K27ac before DOT1Li (Figure 4B-C left plot), suggesting that H3K79me3 may be particularly important for maintaining enhancers with only moderate levels of H3K27ac in an open and active configuration. Taken together, our data suggest that loss of H3K79me3 reduces H3K27ac levels and chromatin accessibility at specific enhancers. This suggests that H3K79me3 may modulate enhancer function at a subset of enhancers to promote transcription.

### Loss of H3K79me perturbs transcription factor binding at KEEs

Our results so far indicate that H3K79me3 is required at KEEs to maintain chromatin accessibility and H3K27ac levels. To explore this mechanism more deeply, we wanted to investigate which transcription factors were enriched at KEEs to determine if transcription factor binding was disrupted by a loss of H3K79me3 (Figure 5A).

**Figure 5.**
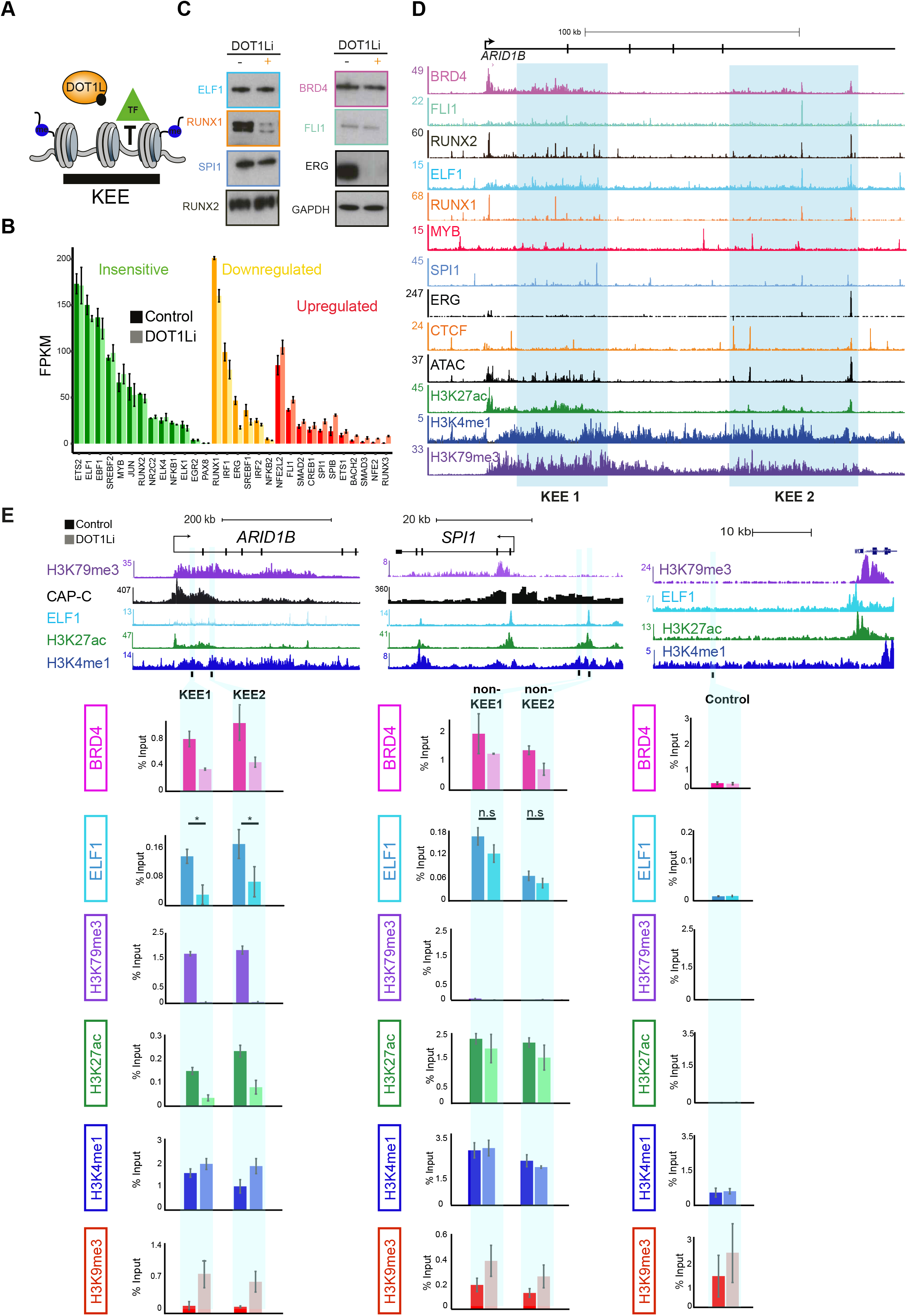
Loss of H3K79me leads to a reduction in transcription factor binding at KEEs (**A**) Schematic showing a gain of H3K9me3, loss of H3K27ac and an antagonism of transcription factor binding at KEEs. (**B**) Expression level of transcription factor motifs enriched at KEEs compared to the rest of the genome. Sorted into categories of transcriptionally insensitive (green), downregulated (orange) and upregulated (red) following DOT1Li. (**C**) Western blot analysis following whole cell extraction of SEM cells in control (-) and DOT1Li (+) conditions. (**D**) ChIP seq tracks for transcription factors in SEM cells at *ARID1B*. Blue bars highlight KEE clusters. (**E**) ChIP qPCR at KEEs and non-KEEs for BRD4, ELF1, H3K79me3, H3K27ac, H3K4me1 and H3K9me3 in control (darker shade) and DOT1Li (lighter shade) conditions. N=3, error bars represent SEM except for ELF1 where N=6. For ELF1, * = p<0.05 using a Mann Whitney U test. See also Figure S5.

To investigate this, we first performed a motif analysis to identify transcription factors that were likely to bind to KEEs. The most highly enriched motifs were filtered by expression level of transcription factors in the leukemia cell line and transcriptional response to DOT1Li, providing three classes of transcription factors; i) insensitive to DOT1Li treatment; ii) downregulated by DOT1Li treatment and iii) upregulated by DOT1Li treatment (Figure 5B). Western blotting was used to validate these results showing that some transcription factors (e.g. RUNX1 and ERG) displayed reduced expression upon DOT1Li treatment, while other transcription factors were unaffected by DOT1Li treatment (Figure 5C). We performed ChIP seq for several transcription factors (FLI1, RUNX1, RUNX2, MYB, SPI1, ERG and ELF1) and found binding at KEEs as well as non-KEEs, with *ARID1B* shown as an example (Figure 5D). To determine whether transcription factor binding is perturbed at KEEs by the loss of H3K79me3, we focused on the transcription factor ELF1, whose protein level is not affected by DOT1Li (Figure 5C). Strikingly, we observed a reduction in ELF1 binding at the KEEs of *JMJD1C, ARID1B* and *BCL11A* (Figure 5E, S5A) but not at *BCL2* or *SPI1* non-KEEs (Figure 5E, S5A). This suggests that some transcription factor binding is specifically reduced at KEEs following loss of H3K79me3.

In order to understand this mechanism further we wanted to identify whether other changes in the chromatin environment at KEEs was observed following the loss of H3K79me3 and subsequent reduction of H3K27ac. Consistent with our ChIP-seq results, H3K27ac levels are reduced at KEEs but not at non-KEEs at several representative loci (Figure 5E, S5A), but no changes in H3K4me1, another enhancer specific modification, were observed (Figure 5E, S5A). BRD4 is a reader of histone acetylation (Dey et al., 2003; Filippakopoulos et al., 2010) and its chromatin binding has been shown to be dependent on H3K79me2/3 and H4K5ac (Gilan et al., 2016). As BRD4 is an important regulatory protein at enhancers and promoters, we asked whether a disruption in BRD4 binding could explain the specific effect of DOT1Li on KEE genes. However, we observed a decrease in BRD4 binding at both KEEs and non-KEEs (Figure 5E, S5A) suggesting that reductions in BRD4 binding is more general and non-specific. This result is consistent with Gilan *et al.*, who saw decreases of BRD4 at the enhancers of some genes that were also regulated by H3K79me3 in the gene body, but also suggests that this effect may be due to effects on transcription elongation that feed back to the enhancer. Similarly, although we observed a modest increase in the repressive modification H3K9me3, this appeared to be a more general effect occurring at both KEEs, non-KEEs and the control region (Figure 5E, S5A), suggesting that this increase is not sufficient to explain specific reductions in transcription factor binding at KEEs. This global change in H3K9me3 can be most likely explained by the upregulation of H3K9me3 methyltransferases such as SETDB2 (Figure S5B) and the downregulation of H3K9me3 specific demethylases such as JMJD1C (Figure S5C) and may also explain the more general disruption to BRD4 binding we observed.

Taking these data together, we propose that the loss of BRD4 and increases in H3K9me3 following DOT1Li cannot explain the specific changes in enhancer characteristics at KEEs compared to non-KEEs. However, there is clear link between H3K79me3-dependent H3K27ac, chromatin accessibility and transcription factor binding at KEEs, consistent with the increased transcriptional sensitivity of KEE genes to DOT1Li. Thus it is likely that it is the specific loss of H3K27ac itself and not the general increase in H3K9me3 that leads to a reduction in chromatin accessibility as well as reduced transcription factor binding following DOT1Li.

### Loss of H3K79me leads to a reduction in KEE-promoter interactions

As we have demonstrated that loss of H3K79me3 at KEEs leads to a reduction in transcription factor binding, we hypothesised this might perturb enhancer function by disrupting enhancer-promoter interactions. To address this, we used the high resolution chromosome conformation technique, Capture-C (Davies et al., 2016). All Capture-C oligos were designed to the promoter of the gene, so that all interactions between the promoter and any potential active enhancers would be captured (Table S2). Overall domain profiles are not affected following DOT1Li, suggesting that domain boundaries are maintained following a loss of H3K79me3 even when there is an associated loss of transcription (Figure 6A-C). However, interactions between KEEs and the promoter within these domains are strongly perturbed by DOT1Li, whereas non-KEE-promoter interactions are unaffected. This is highlighted in the cases of *ARID1B, BCL11A, CDK6, JMJD1C* and *MEF2C* (Figure 6A-B, S6) which show strong KEE-promoter interactions and *BCL2* and *SPI1*, which show strong non-KEE-promoter interactions (Figure 6C, S6). This suggests that H3K79me3 is important for the maintenance of the promoter interaction for KEEs.

To demonstrate this at a wider range of genes, we compared promoter-enhancer interactions at 22 genes in control and DOT1Li conditions. The vast majority of significant differences in promoter interactions were observed with KEEs, with very few promoter-non-KEE interactions affected by DOT1Li (Figure 6D, S6). Most of the promoter-KEE changes were decreases in interactions (Figure 6E, decreases below diagonal line and increases above diagonal line) and were coupled with a downregulation in transcription (Figure 6E, left panel). Although a few non-KEEs did demonstrate a quantifiable reduction in read count of Capture-C signal, these changes were mostly non-significant (Figure 6E, Table S7). Taken together, this suggests that H3K79me3 plays a role in maintaining enhancer-promoter interactions at KEEs and in doing so promotes transcriptional activation of key leukemia gene targets (Figure 7A).

**Figure 6.**
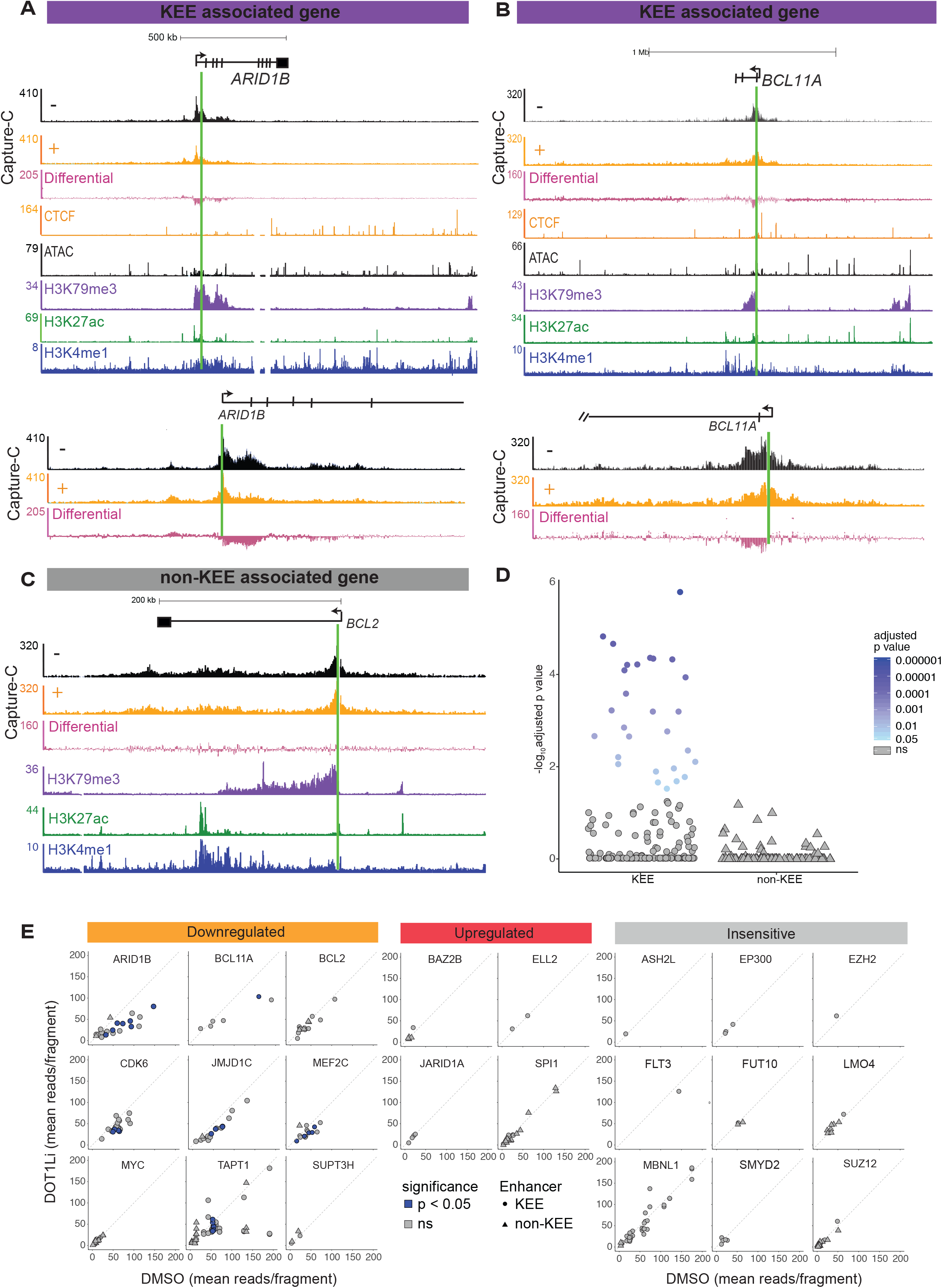
Loss of H3K79me3 leads to a reduction in KEE-promoter interactions (A-C) Capture C (n=3) in control (-, black) and DOT1Li (+, orange) at *ARID1B, BCL11A* and *BCL2*. Differential track demonstrating differences in Capture-C signal (DOT1Li minus control, pink). Capture-C probes designed and captured from the promoter of the gene in all cases (green line). (**D**) Statistical analysis of all Capture-C changes observed at KEEs and non-KEEs. (**E**) Statistical analysis of Capture-C performed in triplicate from the promoter of KEE- and non-KEE genes. X-axis represents read count in control treatment vs read count in DOT1Li condition on Y-axis. Each point represents a KEE (circle) or non-KEE (triangle) that interacts with the gene promoter. Blue points show significant (p<0.05) changes, grey points show non-significant changes. See also Figure S6.

**Figure 7.**
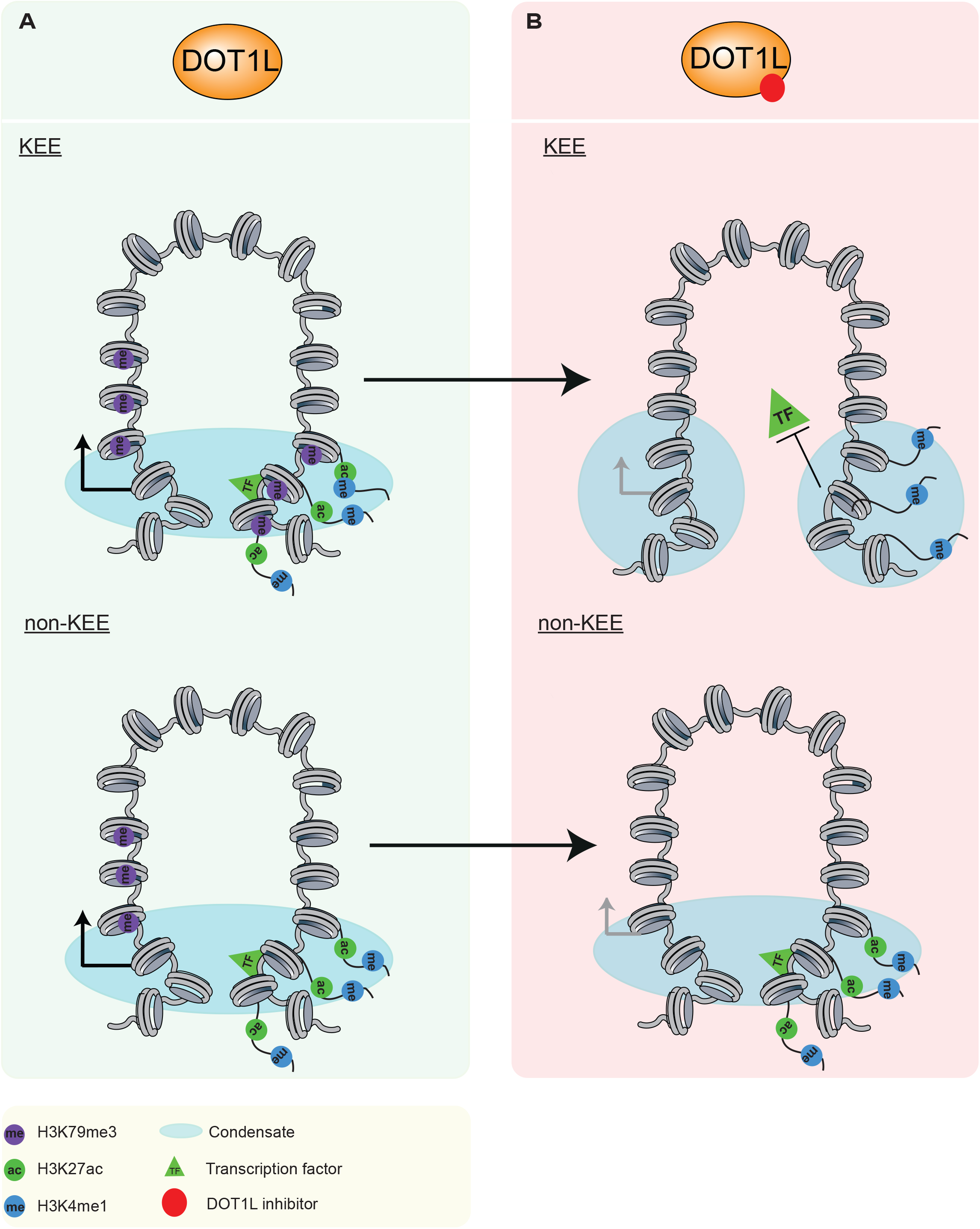
Model of H3K79me3 function at KEEs. (**A**) Schematic diagram representing KEEs (top half) which are marked with H3K79me3 (purple), H3K27ac (green) and H3K4me1 (blue) and bound by specific transcription factors (green triangle). Blue shading represents phase separated condensate between KEE and promoter. Non-KEEs marked with H3K27ac and H3K4me1 only (bottom half). (**B**) Schematic diagrams at KEEs (top half) and non-KEEs (bottom half) following DOT1Li. This shows a loss of the enhancer-promoter condensate and thus loss of interactions upon loss of H3K79me3 at the KEE, but continued maintenance of the enhancer-promoter interactions at the non-KEE even with reduced transcription.

## Discussion

In this study, we demonstrate that H3K79me3 marks a distinct class of enhancers and loss of H3K79me3 impacts enhancer function. We show that a loss of H3K79me3, by treatment with EPZ-5676, leads to a specific reduction of H3K27ac, chromatin accessibility and transcription factor binding at KEEs. Ultimately, this leads to a reduction in KEE-promoter interactions and is coupled with transcriptional downregulation (Figure 7A). This contrasts with our observations at non-KEE regulated genes, such as *BCL2*, where loss of H3K79me3 in the gene body disrupts transcription without impacting enhancer-promoter interactions (Figure 7B).

Past work has suggested that H3K4me1 can control long range enhancer-promoter interactions (Yan et al., 2018), although loss of H3Kme1 alone appears to have very little impact on gene expression (Dorighi et al., 2017). Interestingly, although we see a loss of interactions at KEEs, we do not observe any changes in H3K4me1 suggesting that the role of H3K79me3 in enhancer function is downstream of H3K4me1 deposition and may perhaps indicate that following loss of H3K79me3 and H3K27ac, KEEs revert back to a more poised state (Creygton et al., 2010). We also observed lower levels of H3K27ac at KEEs compared to non-KEEs. This may highlight the importance of H3K79me3 at KEEs which may compensate for the lower acetylation levels. It has been shown that the DOT1L complex members AF9 and ENL can bind to acetylated lysine residues via the YEATS domain (Li et al., 2014, Wan et al., 2017). Therefore, this may provide a positive feedback loop whereby the DOT1L complex is stabilised at KEEs via histone acetylation.

The functional significance of the loss of enhancer-promoter interactions following DOT1Li is a key question which arises from our work. Past observations have demonstrated that one function of H3K79me3 is to inhibit the histone deacetylase SIRT1, and prevent subsequent increases in repressive histone modifications (Chen et al., 2015). This cascade of events would create a more repressive chromatin environment that may disrupt KEE activity potentially by perturbing transcription factor binding (Petruk et al., 2017). However, our work shows that there is a more global increase in H3K9me3 at KEEs and non-KEEs and at the control region, meaning that increases in repressive histone modifications do not explain the specific loss of enhancer-promoter interactions we observe at KEEs.

Instead, the specific loss of H3K27ac may be responsible for the reduction in KEE-promoter interactions that we observe. Recent work has suggested that enhancers may act as hubs for the accumulation of factors that create phase separated condensates generating distinct regulatory domains (Cho et al., 2018; Sabari et al., 2018). Therefore, a loss of H3K79me3 and H3K27ac may be responsible for the disruption of interactions within such a hub so that it can no longer stably promote transcription.

In summary we describe here a distinct set of active enhancers which are dependent upon H3K79me3 to promote transcription. H3K79me3 stabilizes KEE-promoter interactions, maintains H3K27ac and binding of transcription factors, potentially generating phase separated condensates.

## Acknowledgments

T.A.M., L.G., N.T.C., R.T., D.D., I.L were all funded by Medical Research Council (MRC, UK) Molecular Haematology Unit grant MC_UU_12009/6 and MR/M003221/1. D.J.D was supported by a Wellcome Trust Strategic Award (106130/Z/14/Z). The research was also supported by the National Institute for Health Research (NIHR) Oxford Biomedical Research Centre (BRC) Programme.

## Author contributions

Conceptualisation, L.G., T.A.M., N.T.C.; Methodology, L.G., T.A.M., N.T.C., D.J.D., A.M.O., J.R.H.; Software, J.M.T.; Validation, L.G., T.A.M, D.D., Formal Analysis, N.T.C, R.T, J.M.T, E.P.; Investigation, L.G, N.T.C, I.L, D.D, T.A.M.; Data curation, N.T.C, J.M.T.; Writing - original draft, L.G, T.A.M.; Writing - Review and editing, P.V., L.G, N.T.C, J.M.T, A.M.O, D.J.D, J.R.H, T.A.M.; Visualisation, L.G, N.T.C, T.A.M.; Supervision, J.R.H, T.A.M., Funding Acquisition, P.V, J.R.H, T.A.M.

## METHODS

### EPZ-5676 treatment

SEM cells (cultured in IMDM) were seeded at 0.3×10^6^ cells/ml in 50ml of media. Cells were treated with 2μM EPZ-5676 or with 0μM (DMSO only) control. Cells were grown for 7 days, with a change of fresh media and EPZ-5676 at day 3 and 6, where the cells were counted and split to 0.5×10^6^ cells/ml and 0.7×10^6^ cells/ml, respectively. At day 7, EPZ-5676 treated cells were harvested and processed for downstream applications.

### Salt soluble cell extraction and histone acid extraction

A salt soluble cell extraction was performed on 1×10^6^ SEM cells using BC300 (20mM Tris HCl pH 7.5; 20% glycerol; 300mM KCl; 5mM EDTA) + 0.5% NP40 + Protease inhibitor cocktail (Roche). Following this an acid extraction of histone proteins was performed using 0.4M HCl and acetone. Samples were then processed using western blot analysis.

### Western Blot Analysis

Western blot analysis was performed as previously described (Wilkinson et al., 2013).

### Chromatin immunoprecipitation assays

ChIP and ChIP-seq experiments were carried out as previously described (Benito et al., 2015; Kerry et al., 2017; Wilkinson et al., 2013). In brief, fixed samples of up to 1×10^8^ SEM cells were sonicated using a Covaris (Woburn, MA) according to the manufacturer’s recommendations. Ab:chromatin complexes were pulled down using a mixture of magnetic Protein A and Protein G Dynabeads (Life Technologies) and were washed three times with a solution of 50mM Hepes-KOH, pH 7.6, 500mM LiCl, 1mM EDTA, 1% NP-40, and 0.7% Na deoxycholate.

Following a Tris-EDTA wash, samples were eluted, treated with RNase and proteinase K, and purified using a Qiagen PCR purification kit. DNA was then analysed using qRT-PCR, with ChIP samples quantified relative to inputs (Milne et al., 2009) (Table S1 for list of qPCR primers used). For ChIP-seq, DNA libraries were made using the NEBnext ultra DNA library preparation kit for Illumina (Cat no. E7370). Samples were sequenced on a Nextseq 500 using paired end sequencing.

**Table S1:**
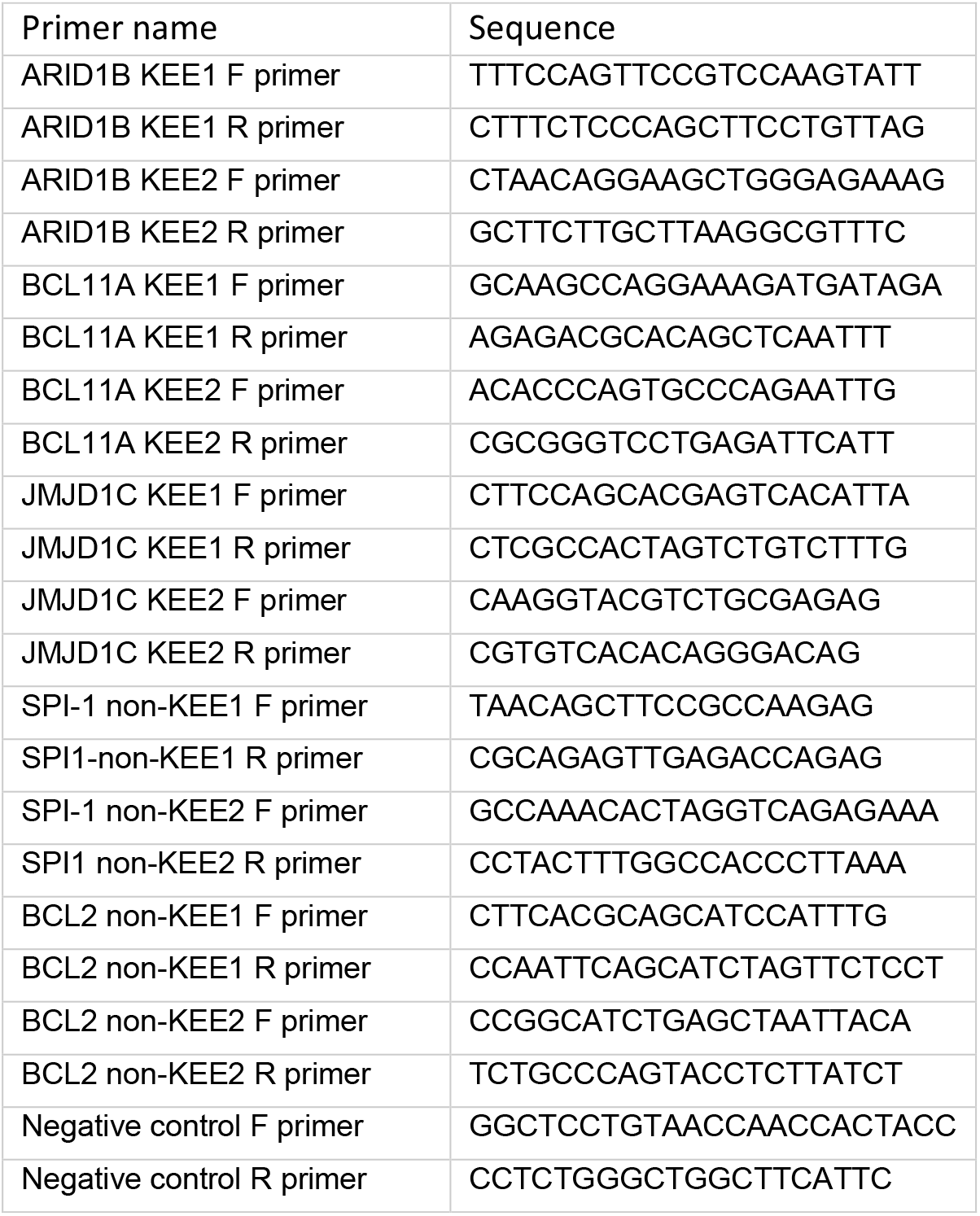
qPCR primer list. Related to Figure 5 and S4

### ChIP-rx sequencing

ChIP-rx sequencing was performed as described in Orlando et al., 2014. Briefly, fixed *Drosophila melanogaster* S2 cells were added at the lysis step of the ChIP to SEM cells at a ratio of 1:4. Following sequencing, reads were mapped to both the hg19 and dm3 genome builds using an in-house pipeline.

### Nascent RNA extraction

Following 7 day EPZ-5676 treatment, 1×10^8^ cells were treated with 500μM 4-thiouridine (4-SU) for 1 hour. Cells were lysed using trizol and RNA was precipitated with ethanol. 4-SU-incorporated RNA was biotinylated by labelling with 1mg/ml Biotin-HPDP for 90 minutes at room temperature. Following chloroform extraction, labelled RNA was separated using magnetic streptavidin beads. Beads were washed using a magnetic μMACS stand before RNA was eluted in two rounds of elution with 100μL 100mM DTT. RNA was purified using a Qiagen RNeasy MinElute kit. Samples were sequenced using a NextSeq 500 and paired-end sequencing.

### Nascent RNA-seq and Gene Expression Analysis

Following QC analysis with the fastQC package (http://www.bioinformatics.babraham.ac.uk/projects/fastqc), reads were aligned using STAR (Dobin et al., 2013) against the human genome assembly (hg19). Reads that were identified as PCR duplicates using Samtools (Li et al., 2009) were discarded. Gene expression levels were quantified as read counts using the featureCounts function (Liao et al., 2014) from the Subread package (Liao et al., 2014) with default parameters. The read counts were used for the identification of global differential gene expression between specified populations using the edgeR package (Robinson et al., 2010). RPKM values were also generated using the edgeR package. Genes were considered differentially expressed between populations if they had an adjusted p-value (FDR) of less than 0.05.

### ATAC-seq

The ATAC seq protocol was adapted from (Buenrostro et al. 2013). 50 000 SEM cells were harvested and washed in PBS and resuspended gently in 50μl cold lysis buffer (10mM Tris-HCl, pH 7.4; 10mM NaCl; 3mM MgCl2; 0.1% IGEPAL CA-630). Cells were spun down immediately at 500xg for 10min at 4°C. The pellet was resuspended in a transposase reaction mix at 37°C for 30 minutes. DNA was purified using a Qiagen MinElute Kit as per manufactures instruction. The DNA fragments were amplified in a PCR reaction and purified using Qiagen PCR clean up kit. Samples were sequenced on a NextSeq 500 using a 75 cycle kit. Statistical analysis of ATAC peaks from five replicates was conducted with Diffbind, using EdgeR. Peaks were considered different between conditions if they had an adjusted p-value (FDR) of less than 0.05.

### Sequence Analysis

For ChIP-seq, ChIP-rx and ATAC-seq, quality control of FASTQ reads, alignment, PCR duplicate filtering, blacklisted region filtering and UCSC data hub generation was performed using an in-house pipeline https://github.com/Hughes-Genome-Group/NGseqBasic/releases. Briefly, the quality of the FASTQ files were checked with fastQC, then mapped using Bowtie against the human genome assembly (hg19). Unmapped reads were trimmed with trim_galore and then mapped again. Short unmapped reads from this step were combined using Flash and then mapped again. PCR duplicates were removed using samtools rmdup, and any read mapping to Duke blacklisted regions (UCSC) were removed using bedtools. Directories of sequence tags (reads) were generated from the sam files using the Homer tool makeTagDirectory. The makeBigWig.pl command was used to generate bigwig files for visualisation in UCSC, normalising tag counts to tags per 10 million. For ChIP-rx, the normalisation factor was adjusted to take into account the ratio of mapped human and Drosophila reads in the bound and input samples.

Peaks were called using the Homer tool findPeaks, with the input track provided for background correction, using the-style histone or-style factor options to call peaks in histone modification or transcription factor/ATAC datasets, respectively. Metagene profiles were generated using the Homer tool annotatePeaks.pl.

### Enhancer State Identification

Enhancer states were called using ChromHMM. Briefly, the genome was subdivided into 200 bp buckets, and each bucket was iteratively assigned to one of 30 states using the following ChIP-seq peak files from SEM cells: H3K4me1, H3K4me3, H3K9ac, H3K27ac, H3K27me3, H3K36me3, H3K79me3. States were then recombined based on the presence/absence of modification peaks. Specifically, KEEs were defined as the presence of H3K4me1, H3K27ac and H3K79me3. Conversely, non-KEEs were defined as the presence of H3K4me1 and H3K27ac in the absence of H3K79me3. Enhancer buckets less than 1 kb apart were merged together, and the merged enhancer was labelled a KEE if a KEE bucket was present. Enhancers were labelled intragenic if they overlapped with a gene’s coordinates, or intergenic if not. Enhancers were assigned to the nearest gene.

### Capture-C

Next-generation Capture-C was performed as previously described (Davies et al., 2016). 2 × 10^7^ SEM cells were assayed. 3C libraries were sonicated to a fragment size of 200 bp and Illumina paired-end sequencing adaptors (New England BioLabs, E6040, E7335 and E7500) were added using Herculase II (Agilent) for the final PCR. Indexing was performed in duplicate to maintain library complexity with libraries pooled after indexing. Capture probes targeting promoters were designed as 120 bp biotinylated DNA oligonucleotides (IDT) using the online CapSequm tool (http://apps.molbiol.ox.ac.uk/CaptureC/cgi-bin/CapSequm.cgi; Hughes 2014) (Table S2, list of biotinylated capture-c oligos). Enrichment was performed with two successive rounds of hybridization, streptavidin bead pulldown (Invitrogen, M270), bead washes (NimbleGen SeqCap EZ) and PCR amplification (Kapa/NimbleGen SeqCap EZ accessory kit v2). The material was sequenced using the Illumina NextSeq platform with 150-bp paired-end reads. Data analysis was performed using an available pipeline (github.com/Hughes-Genome-Group/CCseqBasicF/releases; Davies 2016).

Capture-C interactions between captured promoters and enhancers were quantified for statistical analysis. Peaks outside of the bounds of Capture-C interaction domains (visually determined using UCSC genome browser) and those on trans chromosomes were removed from the analysis. Peaks within 10 kb of the Capture-C probe hybridisation site were also removed. p-values for each peak were calculated by comparing all of the normalised read counts for each DpnII fragment and all replicates within a peak using a paired Mann-Whitney test for the two treatment conditions (DMSO vs. EPZ-5675 2μM).

## QUANTIFICATION AND STATISTICAL ANALYSIS

All statistical analysis was performed using R (v3.3.3) (R Core Team, 2018). Between-group comparisons of continuous variables were performed with the Wilcoxon rank sum test. Contingency table tests were performed with Fisher’s exact test. Exact p-values can be found in Table S6. Capture-C statistics can be found in Table S7.

## DATA AND SOFTWARE AVAILABILITY

High throughput data has been submitted to GEO. Datasets used from previous publications can be found in Table S8.

**Table S8.** Datasets from previous publications. Related to Figures 1–6 and S1–6

| ChIP seq |  |  |
| --- | --- | --- |
| Cell line | Antibody | Accession number |
| SEM | H3K79me3 | GSE83671 |
|  | H3K27ac |  |
|  | H3K4me1 |  |
|  | H3K4me3 |  |
|  | BRD4 |  |
|  | RUNX1 | GSE42075 |
| Nascent RNA seq |  |  |
| Cell line | Treatment | Accession number |
| SEM | Control replicate 1 | GSE83671 |
|  | Control replicate 2 |  |
|  | Control replicate 3 |  |
|  | DOT1Li replicate 1 |  |
|  | DOT1Li replicate 2 |  |
|  | DOT1Li replicate 3 |  |

## Supplementary Information Figure Legends

**Figure S1, related to Figure 1.**
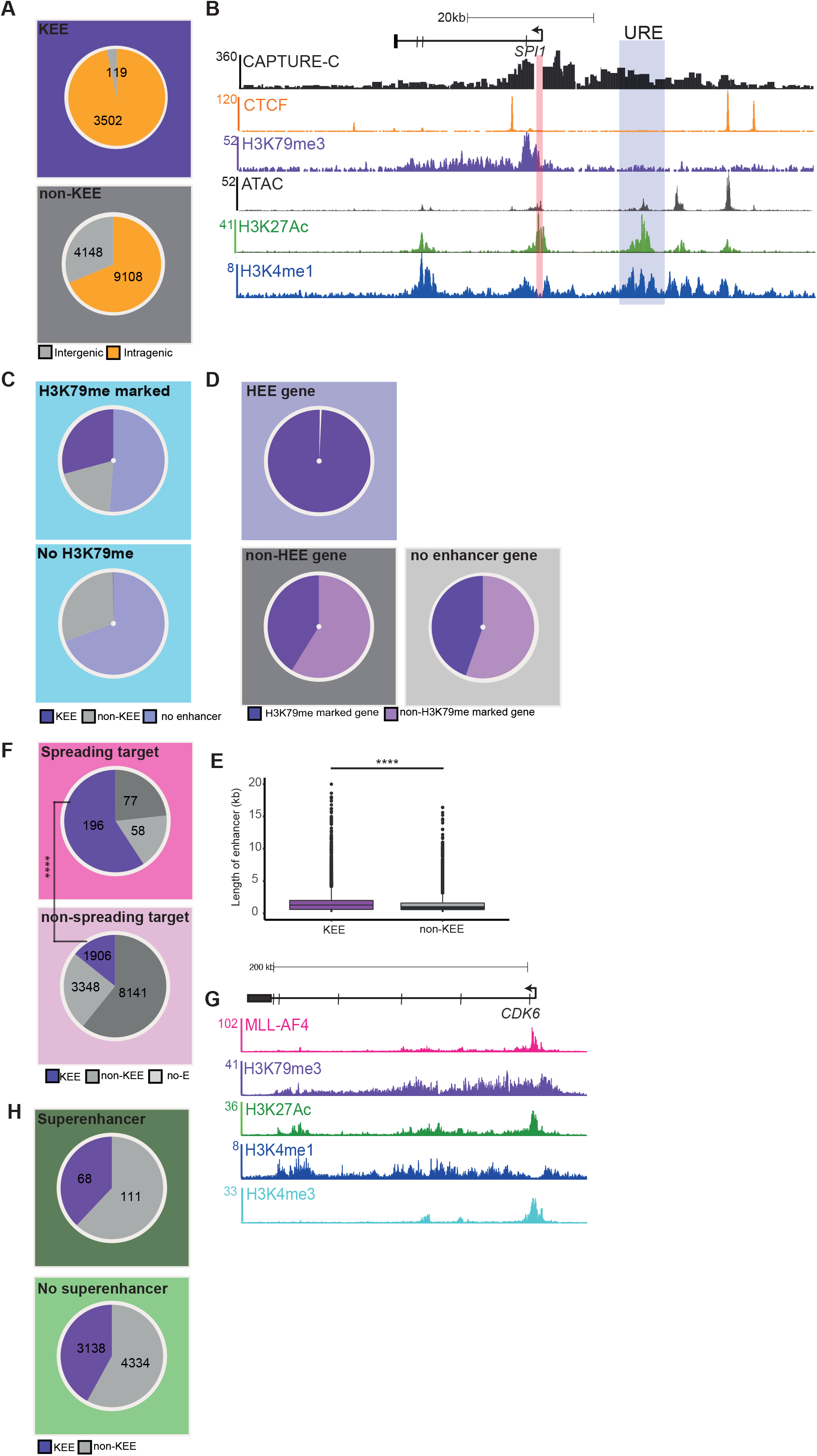
H3K79me3 marks a distinct class of enhancer. (**A**) Proportion of intragenic (orange) and intergenic (grey) KEEs and non-KEEs. (**B**) Example of a gene, *SPI1*, associated with a non-KEE. (**C**) Proportion of H3K79me marked genes or genes with no H3K79me that are associated with a KEE, non-KEE or no enhancer. (**D**) Proportion of KEE genes, non-KEE genes and no enhancer genes which are marked or not marked with H3K79me. (**E**) Distribution of lengths of KEEs (purple) compared to non-KEEs (grey) (**** =p-value <0.0001 using a Mann-Whitney *U* test). (**F**) Proportion of MLL-AF4 spreading genes associated with a KEE-(purple), non-KEE-(grey) or no enhancer gene (light grey) (****=p-value <0.0001, Fishers exact test). (**G**) Example tracks for an MLL-AF4 spreading gene, *CDK6*. (**H**) Proportion of super-enhancers which are also KEEs or non-KEEs.

**Figure S2, related to Figure 2.**
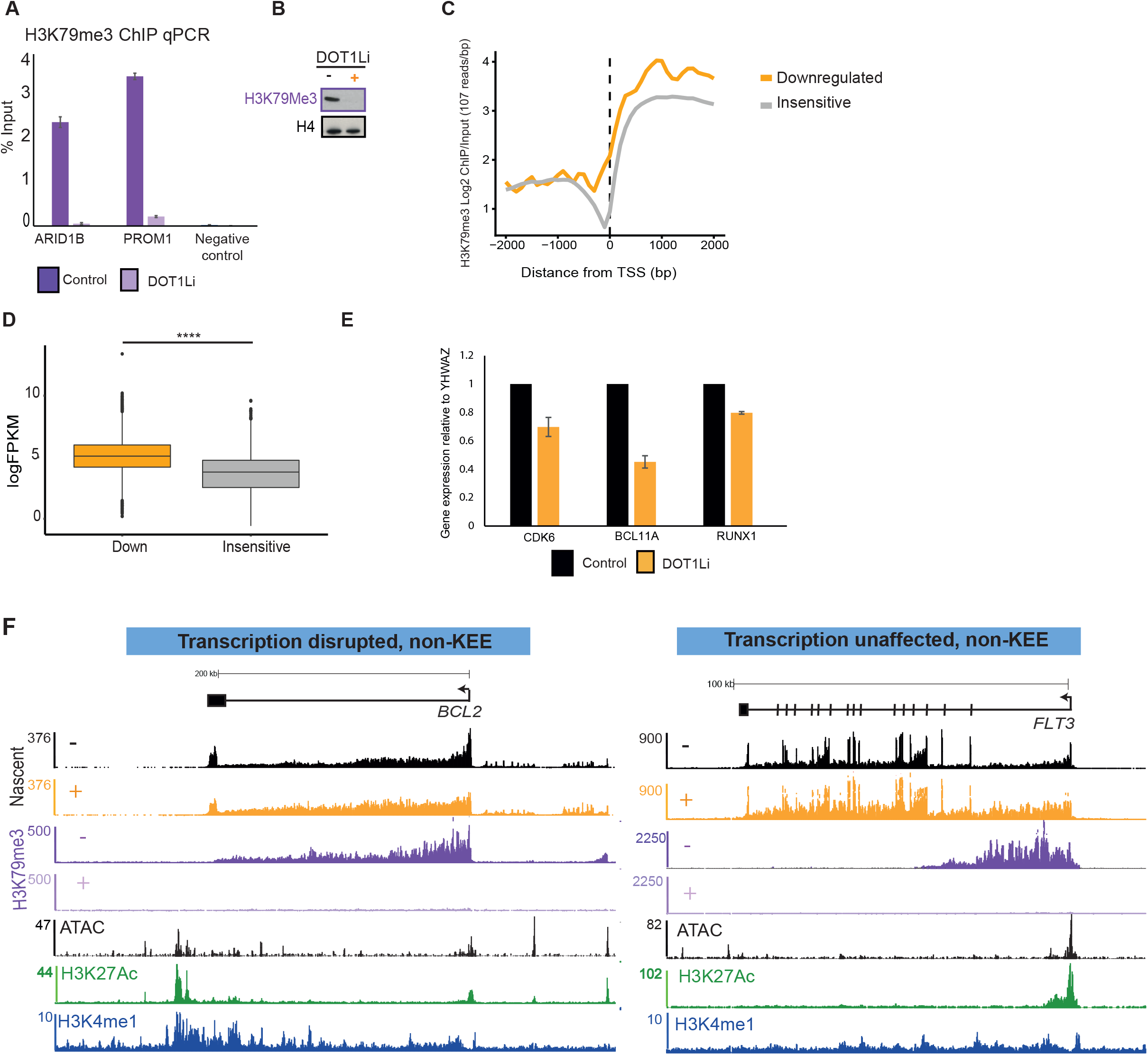
Loss of H3K79me3 leads to a reduction in transcription. (**A**) H3K79me3 ChIP qPCR (n=3) in control (purple) and DOT1Li conditions (lilac) at KEE genes. (**B**) Western blot analysis of H3K79me compared to H4 control in control (purple) and DOT1Li (lilac) conditions. (**C**) Level of H3K79me3 ChIP seq at downregulated (orange) and insensitive genes (grey) following DOT1Li. (**D**) Average expression (LogFPKM) of downregulated (orange) compared to insensitive (grey) (**** =p-value <0.0001 using a Mann-Whitney *U* test). (**E**) qRT-PCR of total RNA (n=3) at KEE genes following DOT 1 Li. (**F**) Example of a gene which demonstrates a reduction in transcription with a non-KEE, *BCL2*. (**G**) Example of a gene which demonstrates no change in transcription and is not associated with an enhancer, *FLT3*.

**Figure S3, related to Figure 3.**
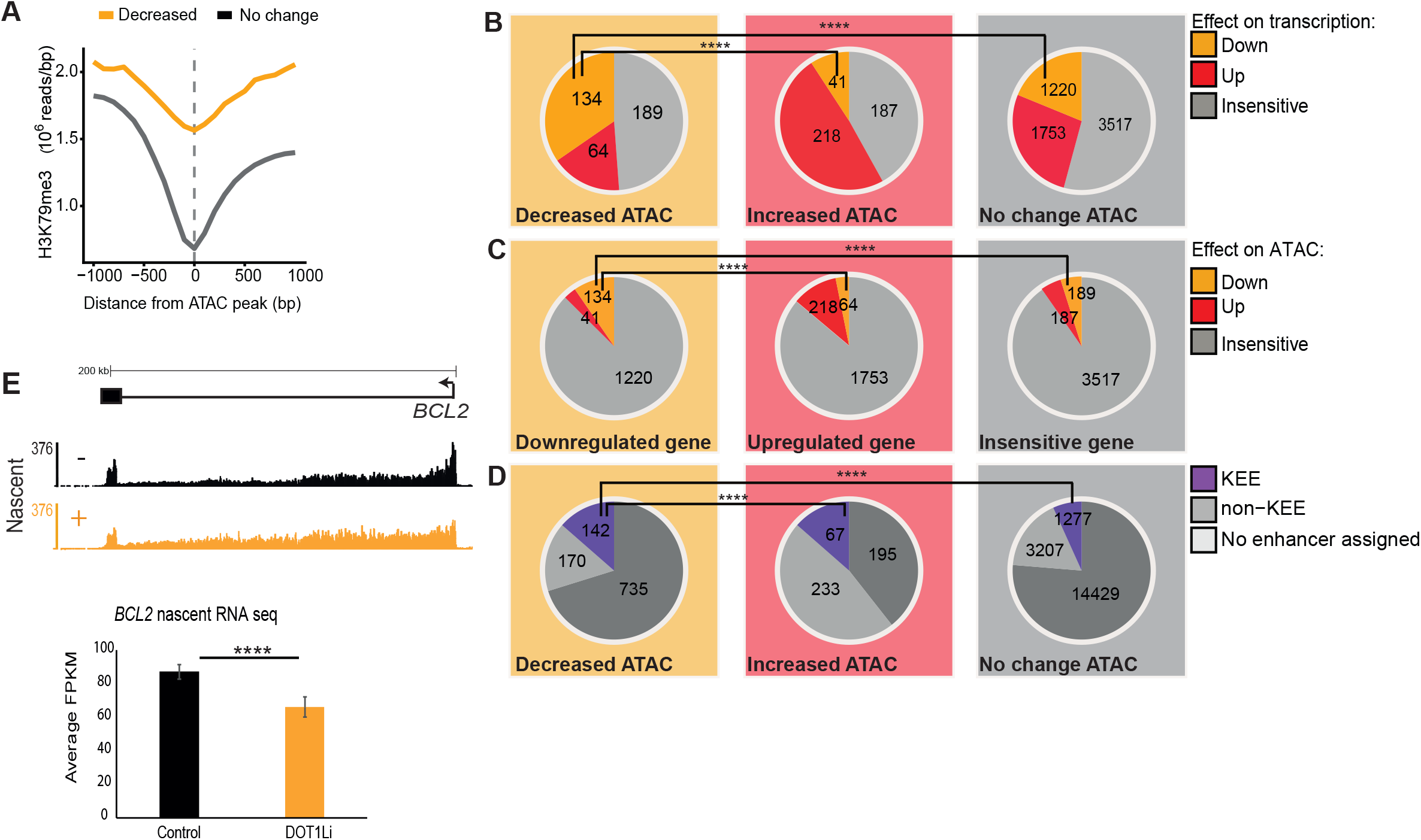
DOT1Li leads to a reduction in chromatin accessibility. (**A**) Metaplot of H3K79me3 ChIP seq signal centered around reduced ATAC peaks and those which demonstrate no change following DOT1Li (**B**) Proportion of ATAC peaks (decrease, increased or no change) which are associated with genes that are downregulated, upregulated or insensitive to DOT1Li (****=p-value <0.0001, Fishers exact test). (**C**) Proportion of genes, either transcriptionally downregulated, upregulated or insensitive that demonstrate changes in ATAC peaks following DOT1Li (****=p-value <0.0001, Fishers exact test). (**D**) Proportion of decreased, increased or no change ATAC peaks which are found within KEE-genes, non-KEEs genes or genes with no enhancer assigned (****=p-value <0.0001, Fishers exact test). (**E**) Nascent RNA seq of BCL2 in control (black) and DOT1Li (orange) conditions (**** = FDR <0.0001).

**Figure S4, related to Figure 4.**
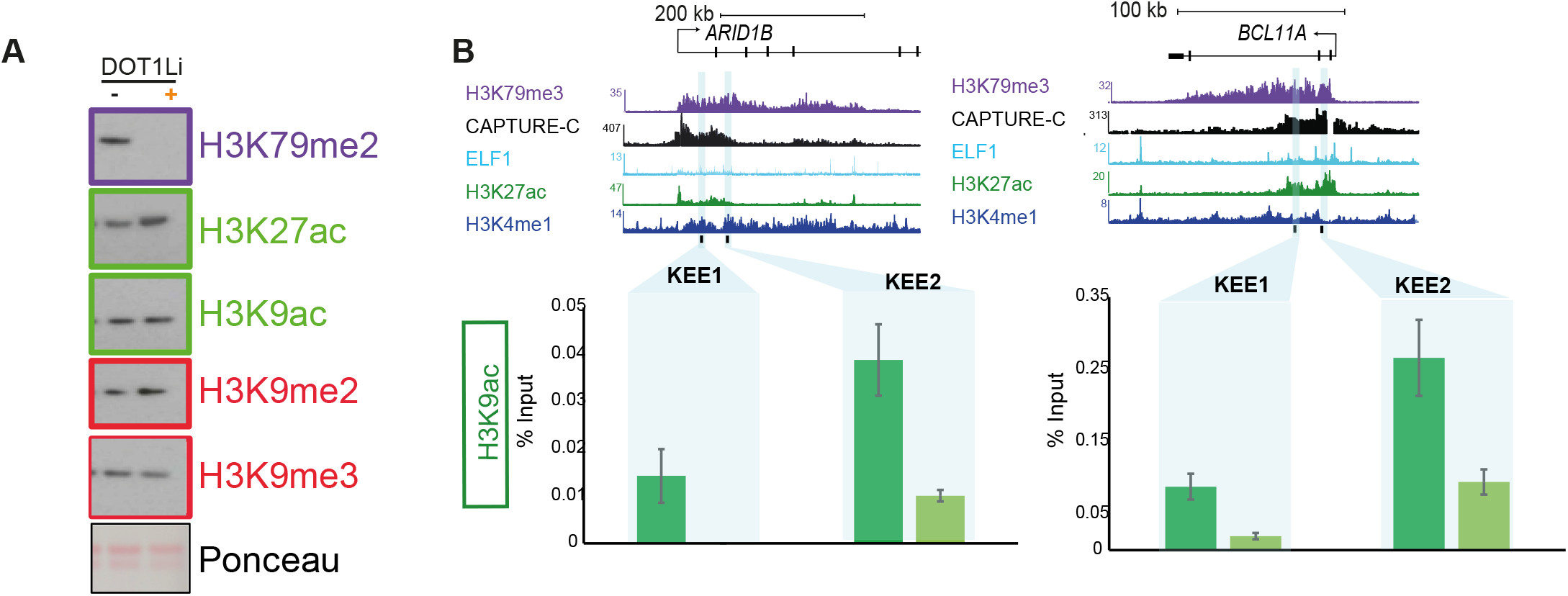
Changes in histone modifications following DOT1Li. (**A**) western blot analysis of acid extracted histones from 1×10^6^ SEM cells in control and DOT1Li conditions (**B**) H3K9ac ChIP qPCR in control (green) and DOT1Li (light green) conditions at *ARID1B* KEEs

**Figure S5, related to Figure 5.**
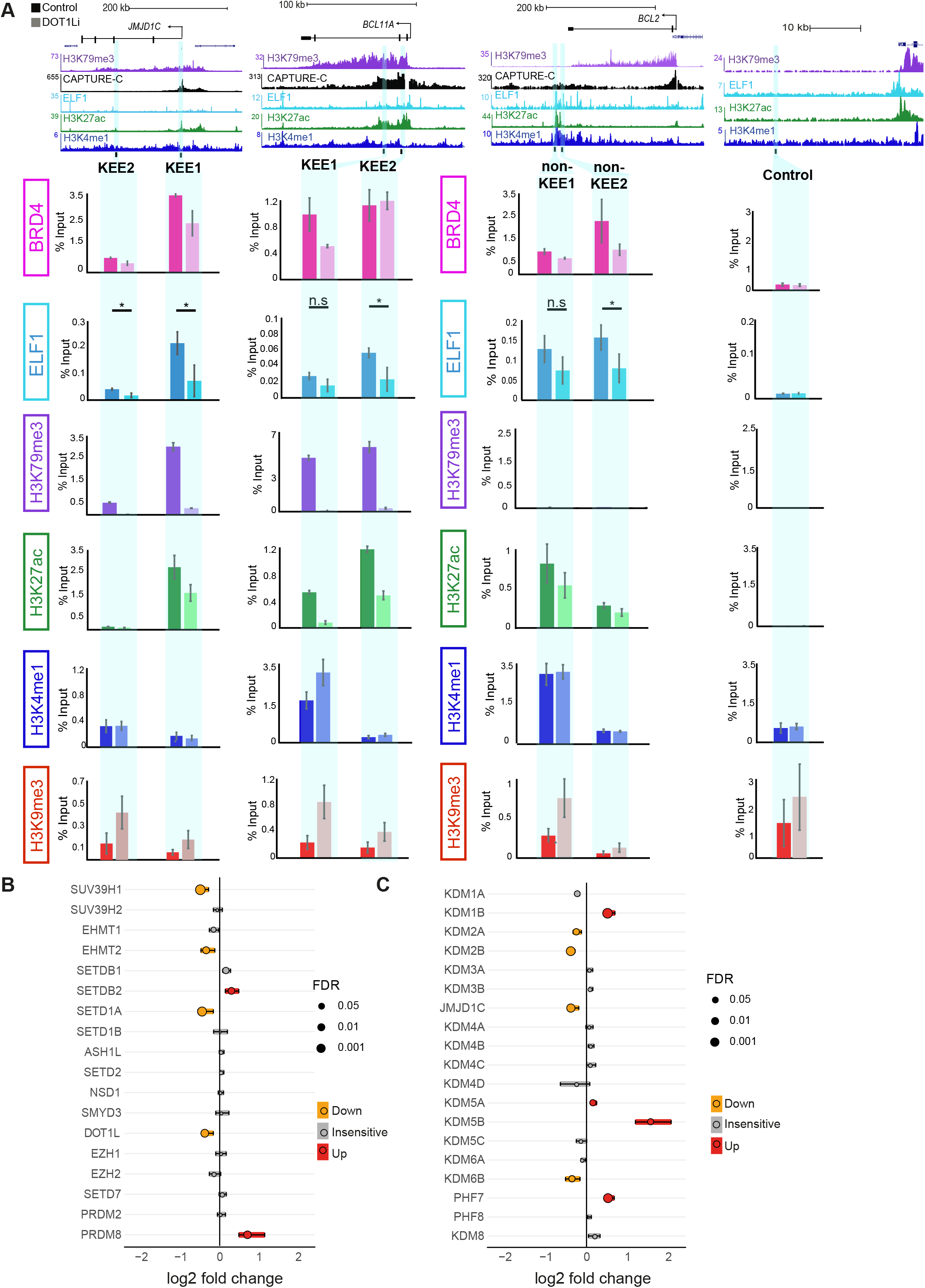
Reduction in transcription factor binding following DOT1Li. (**A**) ChIP qPCR at KEEs and non-KEEs for BRD4, ELF1, H3K79me3, H3K27ac, H3K4me1 and H3K9me3 in control (darker shade) and DOT1Li (lighter shade) conditions. N=3, error bars represent SEM except for ELF1 where N=6. For ELF1 (* = p<0.05 using a Mann Whitney U test). (**B**) Forest plot displaying changes in nascent RNA levels of histone methyltransferases following DOT1Li, either downregulated (orange), insensitive (grey) or upregulated (red). Size of circle represent level of significance of change. Bars represent inter-quartile range (**C**) Changes in nascent RNA levels of histone demethylases following DOT1Li either downregulated (orange), insensitive (grey) or upregulated (red). Size of circle represent level of significance of change. Bars represent inter-quartile range

**Figure S6, related to Figure 6.**
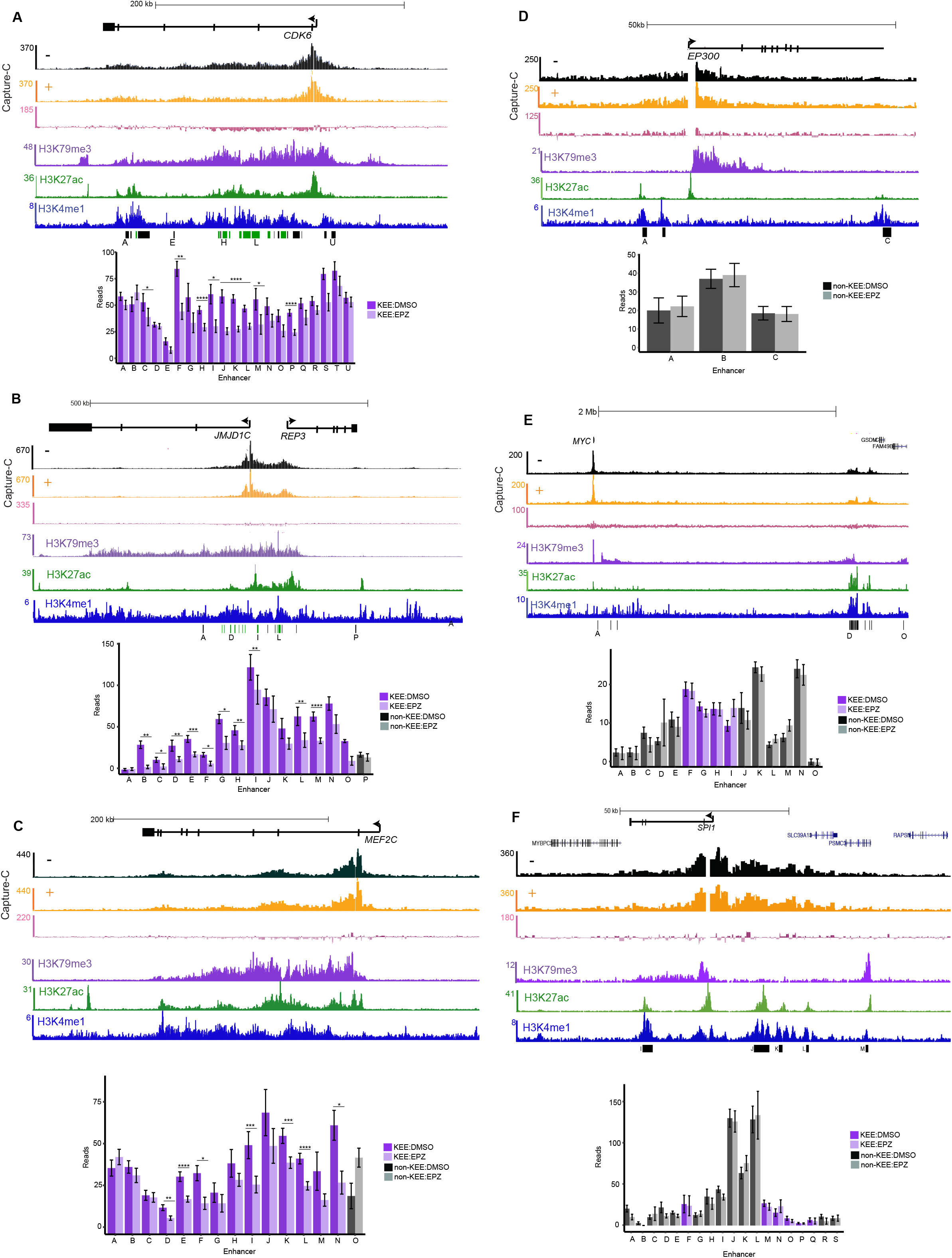
Enhancer-promoter interactions at KEEs and non-KEEs. **(AC)** Capture-C from the promoter of *CDK6, JMJD1C* and *MEF2C* in control (black) and DOT1Li (orange) treatment performed in triplicate. Differential track (pink) demonstrates difference in signal between Control and DOT1Li. Bar charts show statistical significance (Mann Whitney U test) of change in read count between control and DOT1Li conditions and KEEs (purple) and non-KEE (grey) **(D-F)** Capture-C from the promoter of *EP300, SPI1* and *MYC* in control (black) and DOT1Li (orange) treatment performed in triplicate. Differential track (pink) demonstrates difference in signal between Control and DOT1Li. Bar charts show statistical significance (Mann Whitney U test) of change in read count between control and DOT1Li conditions and KEEs (purple) and non-KEE (grey).

